# Revealing synaptic nanostructure distribution through automatic dendritic spine segmentation and single-molecule localization microscopy

**DOI:** 10.1101/2024.07.15.603616

**Authors:** Jiahao Zhang, Rohit Vaidya, Rujuta Pendharkar, Hee Jung Chung, Paul R. Selvin

## Abstract

Relating dendritic spine morphology to synaptic organization in brain tissue is essential for understanding excitatory synaptic transmission and plasticity. Single-molecule localization microscopy (SMLM) offers the spatial precision needed to study the synaptic protein distribution at the nanoscale. However, the widefield setup required for SMLM produces diffraction-limited images with poor contrast and resolution in thick brain slices (> 30 μm), making accurate segmentation of dendritic spines challenging. To overcome this challenge, we developed an automated 3D segmentation approach tailored to this condition by combining two existing machine-learning models. We integrated this strategy with SMLM-based localization of synaptic proteins to map post-synaptic protein PSD-95 within spines at nanoscale resolution. This framework, named ISEPLA (Integrated Spine Extraction and Protein Localization Analysis), revealed a hierarchical organization of synaptic proteins: spines contain multiple nanomodules, each composed of smaller nanodomains. Larger spines contain more nanomodules, and larger nanomodules comprise more nanodomains. Therefore, our method enables precise morphological and molecular analysis under physiologically relevant imaging conditions, providing new insights into the synaptic organization in spines.

## INTRODUCTION

Dendritic spines are the primary sites of excitatory synapses in neurons, providing the morphological foundation for synaptic plasticity.^1–3^ These small protrusions house nanostructures of postsynaptic density (PSD) proteins, which control the transmission and plasticity of the synapse.^4–6^ To understand the interplay between spine morphology and the molecular organization of the PSD proteins in regulating synaptic mechanisms^7–11^, it is critical to simultaneously capture and analyze alterations in overall morphology, i.e., the spine shapes, and the internal nanostructure distribution of PSD proteins.

Obtaining spine structures requires segmentation, which involves delineating the shape of the spine from the dendrites and surrounding tissue.^12–20^ This becomes a significant bottleneck when imaging synaptic proteins using single-molecule localization microscopy (SMLM), a category of super-resolution techniques that can reveal the nanoscale synaptic architectures.^21–24^ Despite the ability of SMLM to map the spatial locations of these nanostructures, these structures cannot be directly attributed to individual dendritic spines without spine segmentation. Therefore, it is necessary to first take a diffraction-limited image of the neuron, followed by SMLM imaging of the synaptic proteins in the same field-of-view (FOV). Because these experiments are typically conducted on dissociated neuronal cultures, the spines from the diffraction-limited image are well-resolved for segmentation. However, there is a growing shift toward applying SMLM to the more physiologically relevant brain tissue.^25–28^ In these thick brain slices, the requirement of a widefield setup (as opposed to a confocal setup) for SMLM experiments results in diffraction-limited images with a low signal-to-background ratio (SBR) and limited resolution. Existing dendritic spine segmentation methods often struggle in these challenging conditions.^12–20^As a result, human annotation is generally required for analysis of SMLM data of synaptic proteins on neurons, which not only introduces potential biases but also becomes labor-intensive for quantitative studies. Hence, current analysis methods lack the capability to explore the connections between the spine structure and the nanoscale organization of synaptic proteins in physiological brain samples.

To address this challenge, we developed a new method called ISEPLA (Integrated Spine Extraction and Protein Localization Analysis), enabling combined analysis of spine structure and synaptic protein distribution. First, to obtain reliable spine segmentations in low SBR and resolution images, ISEPLA combines two open-source machine-learning models, DeepD3 and Stardist. DeepD3 is designed for spine detection and predicts the likelihood of each sub-diffraction-limited pixel belonging to a spine.^29^ The resulting prediction value image is thresholded and used to create a mask on the original fluorescence image, isolating fluorescence intensities in regions with higher prediction values. Stardist, originally developed for nuclei segmentation, then processes these masked intensities to accurately detect and segment the spines.^30^ Separating the segmentation step via Stardist from the detection step via DeepD3, while leveraging fluorescence intensity in both models, improves the accuracy of spine region identification compared to using DeepD3 alone. This segmentation approach can reliably segment spines in 3D and resolve overlapping spines, whereas the traditional segmentation using a watershed algorithm fails.^31^ For benchmarking, we validated the performance of our method for dendritic spine segmentation compared to other approaches using super-resolution imaging as the reference.

On the next step, ISEPLA performs SMLM data analysis of proteins integrated with the spine segmentation results. The custom-written analysis clusters protein localizations to nanosized compartments, which are then assigned to individual spine or dendrite segmentations. Thus, ISEPLA provides an integrative analysis of spine segmentation and SMLM point analysis. In our application, we imaged the PSD marker protein, PSD-95, using Stochastic Optical Reconstruction Microscopy (STORM), a main form of SMLM.^32–34^ PSD95 is known to anchor AMPA and NMDA-type glutamate receptors at excitatory synapses, playing a major role in the maturation, stabilization, and plasticity of excitatory synapses.^35–37^

Using ISEPLA, we gained new insights into understanding the nanoscale organization of PSD-95 inside spines. Previous works have shown that PSDs form sub-diffraction-limited assemblies named “nanodomains” or “nanomodules”, although the distinction between these two categories has remained unclear.^38,39^ In our analysis, we found that while the number of PSD-95 nanomodules increases with spine volume, the average size of the nanomodules does not. Leveraging the high resolution of SMLM, we also resolved the distribution of PSD-95 nanodomains, which have been identified separately in SMLM studies but are below the resolution limit of STED imaging.^40,41^ These nanodomains are known to align with presynaptic release sites via trans-synaptic nanocolumns, regulating synaptic transmission and plasticity.^42^ Importantly, using the quantitative analysis power of ISEPLA, we distinguished nanodomains and nanomodules as different synaptic nanostructures inside spines. A synapse is composed of a single or multiple nanomodules, where each contains highly concentrated regions of nanomodules. We also found that the number of nanodomains scales with the size of the nanomodules, while the ratio of nanodomains to nanomodules remains constant with the spine size.

Therefore, by uncovering the hierarchical organization between dendrite spines and synaptic nanostructures, we showed the power of integrating protein localization data with spine segmentation using ISEPLA.

## RESULTS

### Automatic 3D dendritic spine segmentation for low signal-to-noise and resolution fluorescence images

Here, we describe our automatic 3D dendritic spine segmentation approach. We combined two existing open-source machine-learning models, DeepD3 and Stardist. DeepD3 is an open framework developed for the automated quantification of the number of dendritic spines. It is trained on human-annotated data and has been demonstrated to work for one-photon confocal and ex vivo and in vivo two-photon images taken for different fluorescent labels, species, and brain regions.^29^ However, while it can accurately predict the locations and 2D shapes of spines, it is not focused on deriving the precise segmentation of spines in 3D. For our widefield fluorescence z-stack images, we found that the 3D segmentation function from DeepD3 fails to capture its shape continuously across the z-direction. The 2D shapes of spines can also change drastically across z-frames, despite being similar-looking to the eye. In addition, spines can appear and disappear for a few frames before coming back, which disrupts capturing their shape in 3D.

To tackle this problem, we first attempted to put a threshold on the DeepD3 prediction values and then segment individual spines on the masked binary image using Watershed, a conventional segmentation method.^31^ However, we found that using a low threshold creates oversized spines or creates artificial spines. On the other hand, using a high threshold creates gaps or distorted 2D shapes of spines or misses detection of spines. Moreover, because the Watershed method uses binary images, which do not consider the distribution of fluorescence intensities inside spines, it fails to resolve spines that overlap in space.

Therefore, we needed a way to reliably segment the shapes of spines across z-frames. By noticing the similarity between the geometric shapes of spines and cell nuclei, both of which can be globular or non-globular, we looked to Stardist, an open-source model developed for nuclei segmentation.^30^ By creating segmentations based on the fluorescent intensity distribution rather than a binary image, we found that Stardist can accurately capture the shapes of spines across z-frames. In addition, it can consistently resolve spines that overlap in multiple z-frames. Finally, we connect the segmented 2D shapes across z-frames of a single spine based on their spatial match to create a single 3D segmentation. Thus, our approach uses DeepD3 to first identify regions containing spines in the image, followed by Stardist to segment the spine shapes inside these regions.

The diagram of the workflow for creating 3D segmentation of spines is shown in Figure 1A. After obtaining the fluorescent z-stack of the neuron (step 1), background subtraction is applied to reduce background noise in the image (step 2). Then, for each frame of the z-stack, DeepD3 is used to assign a prediction value each pixel for its likelihood to be part of a spine (step 3). A binary mask is then created by using a prediction value threshold, which is set relatively low, so pixels neighboring the spine regions are included (step 4). This binary mask is mapped back to the original fluorescence image (step 5), and the fluorescence intensity information within the masked regions is extracted (step 6). The intensities inside these regions are used by Stardist to perform 2D segmentation of the spines in each frame (step 7). Finally, the 2D segmentations across different z-axis frames are connected based on spatial overlap to create the 3D segmentation (step 8).

**Figure 1.**
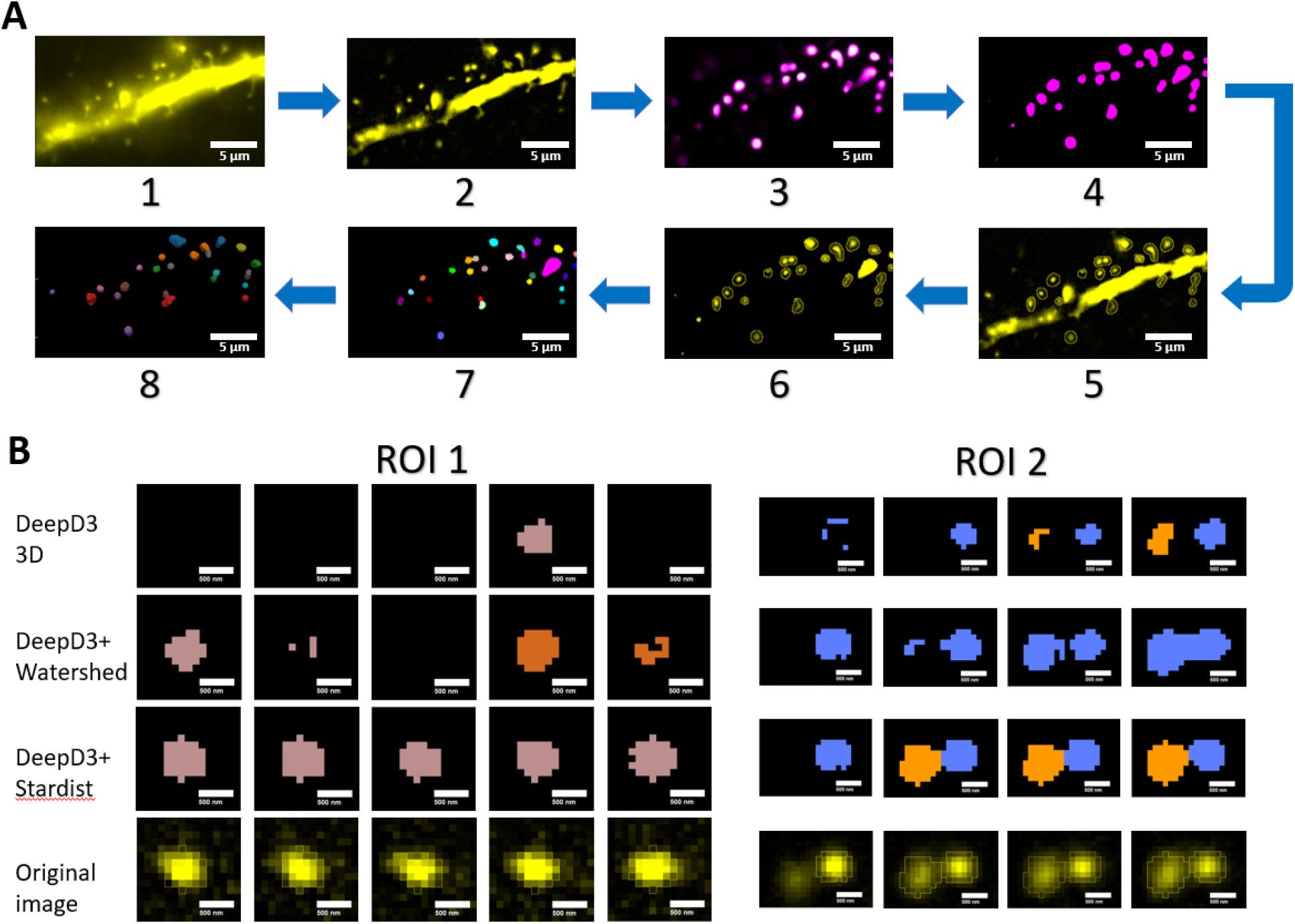
Details of the automatic 3D dendritic spine segmentation approach. (A) Workflow of the approach. 1, The starting z-stack (yellow represents the fluorescence intensity). 2, Subtracting the background. 3, Applying DeepD3 to assign prediction values to the pixels of its likelihood to be part of a spine (magenta to white represents lower to higher likelihood). 4, Creating a binary mask from a low prediction value threshold. 5, Applying the mask back to the original z-stack. 6, Extracting the fluorescent intensity from the masked regions. 7, Applying Stardist to perform 2D segmentation of the spines in each frame (different colors represent differently segmented spines). 8, Connecting the 2D segmentation across each frame to create the 3D segmentation. Scale bar: 5 μm. (B) Example ROIs showing comparison of dendritic spine segmentation results across z-frames for DeepD3 3D segmentation function, DeepD3+Watershed, and DeepD3+Stardist. Scale bar: 500 nm.

Using both DeepD3 and Stardist (DeepD3+Stardist), we can reliably segment single spines in 3D compared to other approaches, with example ROIs in Figure 1B. As shown in ROI 1, the DeepD3 3D segmentation function misses the spine for most of the z-frames despite its presence in the original z-stack. While DeepD3 and Watershed (DeepD3+Watershed) found the spine more frequently, the implementation of a prediction value cutoff still resulted in frames with few or zero pixel segmentation, despite no significant spine shape changes in the original z-stack. The 2D segmentations for z-frames before and after these “gaps” were hence disconnected, resulting in the spine being split into two separate spines. The application of DeepD3+Stardist, on the other hand, segmented the spine consistently across the z-frames that matched with the original z-stack.

The accuracy and consistency of DeepD3+Stardist segmentation were retained even for the case with two spatially overlapping spines. As shown in ROI 2, the DeepD3 3D segmentation function generated distorted shapes with few pixels for some z-frames, which did not match the original image. Furthermore, the DeepD3+Watershed failed to segment the two overlapping spines due to the limitation of the Watershed algorithm, merging the two spines into a single spine with a contorted 3D shape. In contrast, the DeepD3+Stardist not only split the two spines reliably but also matched their shapes precisely to the original z-stack.

### Validation of the dendritic spine segmentation approach with super-resolution images

We validated our approach for dendritic spine segmentation by comparing its match with manual segmentation of super-resolution reconstructions of the same FOV, using STORM or structured illumination microscopy (SIM)^43^ imaging. We took diffraction-limited z-stacks of the dendritic segments of neurons expressing YFP (Yellow Fluorescent Protein) in the hippocampus or cortex of 30 μm thick brain slices (Supplementary Figure 1A-1). The use of Thy1-YFP transgenic mice enabled only a small fraction of neurons to express YFP so their morphology could be examined individually. While SIM imaged YFP directly, STORM imaged the YFP-labeled neurons with an anti-YFP nanobody conjugated to a STORM-compatible dye (Supplementary Figure 1A-2).

For comparison within the different segmentation approaches, we created maximum projections of 40-frame z-stacks of STORM or SIM images and manually segmented them (Supplementary Figure 1A-3). These manual segmentations of super-resolution images were used as ground truth to compare the four segmentation alternatives: our approach using DeepD3+Stardist (Supplementary Figure 1A-4), DeepD3+Watershed (Supplementary Figure 1A-5), the 3D segmentation function in DeepD3 (Supplementary Figure 1A-6), and the commonly used commercial software Imaris (Supplementary Figure 1A-7).

We first compared the segmented spine shapes using intersection over union (IoU), which measures the fraction of pixels matching the ground truth. We found an IoU of 0.38 with STORM reconstruction and 0.34 with SIM reconstruction for DeepD3+Stardist. The original DeepD3 method, compared against human annotation on the same two-photon images, found an IoU of around 0.5.^29^ Given the difference in SNR and resolution of widefield fluorescence versus STORM or SIM, we considered our IoU value to be satisfactory.

While IoU of the projection image quantifies the percentage of pixels correctly identified as part of a spine in 2D, it does not account for consistent segmentation across z-frames. To accurately obtain the shape of a spine, the spine should be consistently detected and segmented in every z-frame along its volume, and overlapping spines also need to be separated. Thus, we also compared the percentage of correctly detected spines, referred to as accuracy. A segmented spine is correctly detected if it is also present in the ground truth. It is incorrectly detected if present only in the ground truth (missed detection) or only in the segmentation from the methods (false detection).

The comparison of our approach with DeepD3+Stardist to other approaches is shown in Supplementary Figure 1B-C. Compared to DeepD3+Watershed, DeepD3+Stardist yielded marginally better IoU, but significantly higher accuracy due to the consistent segmentation across z-frames. Compared to either the DeepD3 program 3D segmentation or Imaris, DeepD3+Stardist yielded significantly improved IoU and accuracy. Therefore, we found significant improvements of DeepD3+Stardist in dendritic spine segmentation for widefield diffraction-limited z-stack images of thick brain slices.

### Application to analysis of STORM localization of synaptic proteins inside dendritic spines

Next, we applied our spine segmentation approach to STORM imaging of post-synaptic density proteins to analyze their distribution *within* each spine. We labeled 30 μm thick brain slices with a nanobody against PSD-95 conjugated to a fluorescent dye. For imaging, we first took a diffraction-limited z-stack of YFP dendritic segments in the hippocampus CA1 region of the brain. Then, we performed STORM imaging of PSD-95 in the same FOV and reconstruction by the in-situ PSF retrieval (INSPR) program.^44^ After using DeepD3+Stardist to segment the spines from the z-stack, we created an automatic analysis to examine and quantify the nanoscale distribution of PSD-95 from its STORM localizations associated with spines.

Figure 2A shows an example image of the STORM reconstruction of PSD-95 overlaid with the Thy1-YFP neuron, along with the zoom-in views of two ROIs containing a single spine.

**Figure 2.**
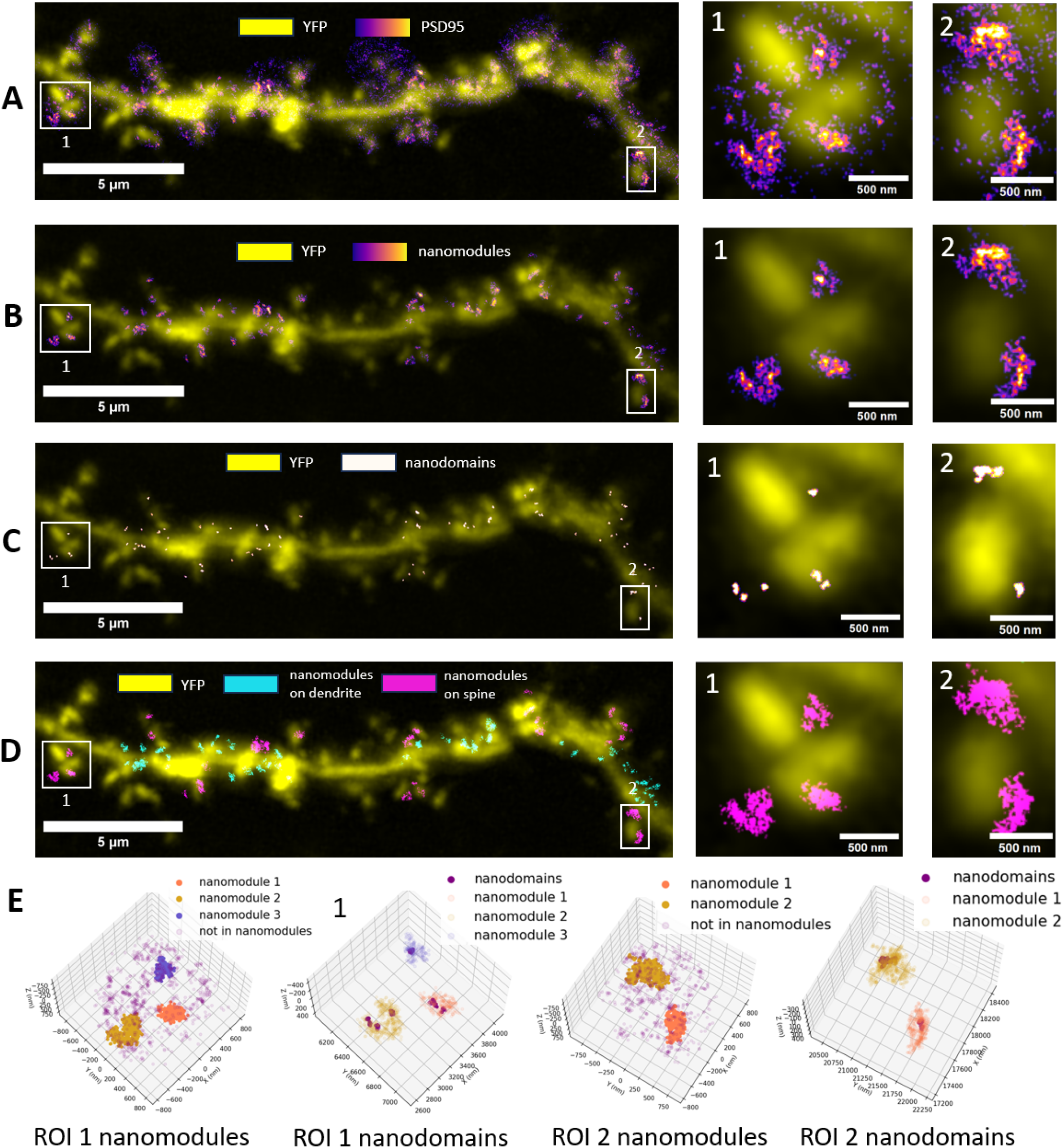
Spine-specific analysis of STORM imaging of synaptic protein PSD-95. (A) STORM reconstruction of PSD-95 (increasing intensity color-coded from purple to bright orange) overlaid on diffraction-limited YFP image (yellow). Two example ROIs contain a single spine. (B) Nanomodules of PSD-95. The same two ROIs and their scatterplots of SMLM localizations. (C) Nanodomains of PSD-95. The same two ROIs and their scatterplots of SMLM localizations. (D) Nanomodules of PSD-95 assigned to individual spines (magenta) or dendrites (cyan).

Localizations over the FOV were assigned to individual spines and dendrites based on the distances to these structures. Figures 2B and 2C show the distribution of PSD-95 nanomodules and nanodomains, respectively, over the neuron. Figure 2D shows nanomodules assigned to individual spines versus the dendrite of the neuron. We confirmed the observation of PSD-95 nanostructures by clustering analysis of localizations based on spatial density by DBSCAN.^45^ The analysis verified the existence of multiple spatially distinct clusters inside a single spine (Figure 2E). These synaptic clusters match the shapes and distributions of synaptic nanomodules found in the 2D STED study.^38^ Here, the individual clusters, or nanomodules, have an average 3D volume of 0.023 μm^3^, consistent with the previously reported area of around 0.1 μm^2^ in 2D by STED. (Assuming a uniform sphere, a volume of 0.023 μm^3^, as determined here, corresponds to a radius of 176 nm and a cross-section area of 0.098 μm^2^, or ∼0.1 μm^2^).^38^

Each nanomodule also has an inhomogeneous density of PSD-95, containing nanoscale high-density regions that match synaptic nanodomains frequently focused on in SMLM studies. To quantitatively confirm this observation, we performed a secondary DBSCAN clustering analysis on localizations belonging to nanomodules. To account for the difference in density of PSD-95 among nanomodules, the density requirement for nanodomains clustering analysis is set to the average density inside nanomodules + 1.5 times the standard deviation. This allowed us to identify the composite nanodomains of nanomodules.

As STED imaging could identify nanomodules inside spines but lacks the spatial resolution to resolve synaptic nanodomains, the relationship between these two terms had not been clearly understood. Past studies mostly focused on either of the two nanostructures, depending on the imaging method and biological question. By combining spine segmentation with STORM clustering analysis, our results suggest that nanodomains are high-density regions inside each synaptic nanomodule.

### Quantification of PSD-95 nanomodules and nanodomains distribution within spines

The spine segmentation and SMLM analysis allowed us to quantify the distribution of PSD-95 nanomodules within the spines, including how their number correlates with spine size. Two-thirds of the spines contained no nanomodules (Figure 3A). Spines containing higher numbers of nanomodules occur less frequently (Figure 3A). By examining the distribution of spine sizes, we found that slightly over half of the spines without nanomodules were small in size (defined as volume < 0.2 μm^3^). In contrast, one-third of spines with nanomodules were small (Figure 3B). Similarly, spines with nanomodules had twice the fraction of large spines (defined as volume greater than 0.6 μm^3^) as spines without nanomodules (29% vs 15%) (Figure 3B). Furthermore, the spine size increased with the number of nanomodules contained within the spine (Figure 3C). This trend is in agreement with the previous STED microscopy study and the general understanding that larger spines tend to be more mature.

**Figure 3.**
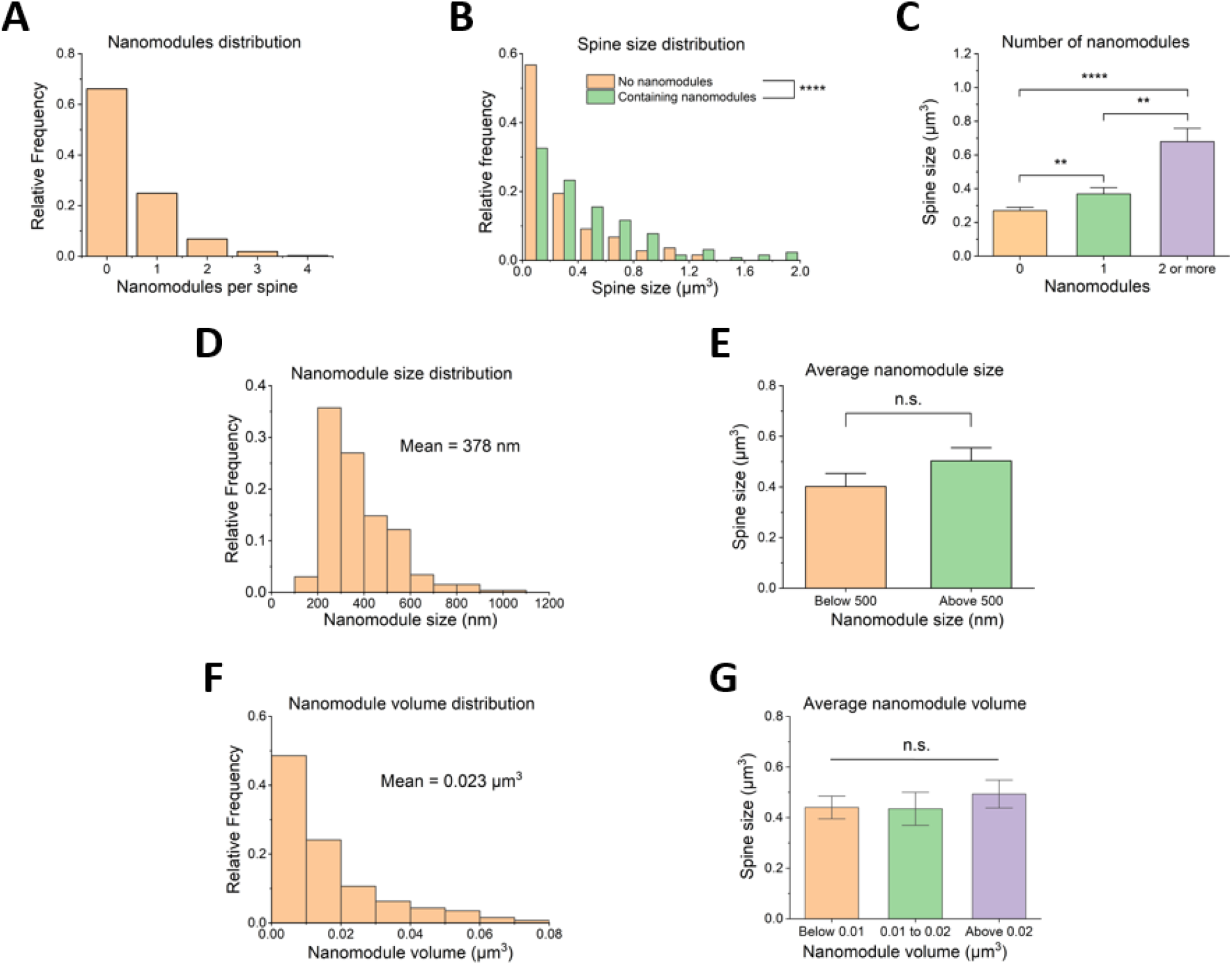
Quantification of nanomodule distribution in spines. (A) Frequency distribution of the number of nanomodules in spines. (B) Frequency distribution of spine sizes, separately for spines containing no nanomodules and for spines containing nanomodules. (C) Spine size versus the number of nanomodules in the spine. (D) Frequency distribution of nanomodule size. (E Spine size versus the average nanomodule size in the spine. (F) Frequency distribution of nanomodule volume. (G) Spine size versus the average nanomodule volume in the spine.

With the distribution of STORM localizations in 3D, we also determined how the sizes/volumes of nanomodules change with spine sizes. The size of each nanomodule is defined as twice the average distance from each localization in the nanomodule to its center. For example, this size will be 0.75 times the sphere radius for a uniformly distributed sphere. Similar to the spine size distribution, nanomodules with increasing sizes occur less frequently, with an average size of 378 nm (Figure 3D). We found that the spine size has no significant dependence on the average size of its nanomodules (Figure 3E). As a second measure of nanomodule dimension, we also examined nanomodule volume, which gives more 3D information, although the specific value is less intuitive than nanomodule radius. Nanomodule volume is found by the volume of the concave hull, i.e., the boundary, built from localizations inside the nanomodule. The fraction of nanomodules similarly decreased with increasing volumes, with an average volume of 0.023 μm^3^ (Figure 3F). In agreement with the trend for nanomodule size, the spine size had *no* significant dependence on the average volume of its nanomodules (Figure 3G). This indicates that larger spines tend to have more nanomodules instead of having larger individual nanomodules.

Significance values are determined by Kolmogorov–Smirnov tests (mean 1 ≠ mean 2) when comparing two cases (B and E) and Kruskal–Wallis ANOVA when comparing more than two cases (C and G) as the data does not follow a normal distribution : p > 0.05, * : p < 0.05, ** : p < 0.01, *** : p < 0.001, ****: p < 0.0001. Data is obtained from 4 independent experiments (from 3 mice), dendritic segments of 12 neurons, and a total of 381 spines. Error bars represent the standard error of the mean.

The application of our segmentation approach and STORM imaging enabled us to examine the statistics of nanodomain distributions inside the nanomodules of the spines. Most nanomodules contained one (45%) or two (26%) nanodomains, with a smaller fraction containing zero (9%) or three (11%) nanodomains, or more nanodomains (9%) (Figure 4A). Nanomodules containing more nanodomains had larger sizes (Figure 4B) or larger volumes (Figure 4C), except for nanomodules with zero nanodomains. Additionally, by calculating the ratio of the number of nanodomains to the number of nanomodules in each spine, we found that this ratio did not significantly depend on the spine size (Figure 4D). This is consistent with our previous finding that the spine size did not depend on the average size of its nanomodules (Figure 3E and 3G), since larger nanomodules contained more nanodomains (Figure 4B and 4C). These findings also predict that the total number of nanodomains would increase with spine size due to an increase in the number of nanomodules. Indeed, we found that spines containing two to three nanodomains were on average larger than spines containing zero to one nanodomain (Figure 4E). Lastly, using the same calculation done for nanomodules, we found the average size of nanodomains to be 66 nm (Figure 4F) and the average volume to be 8.0 × 10^−5^ μm^3^ (Figure 4G).

**Figure 4.**
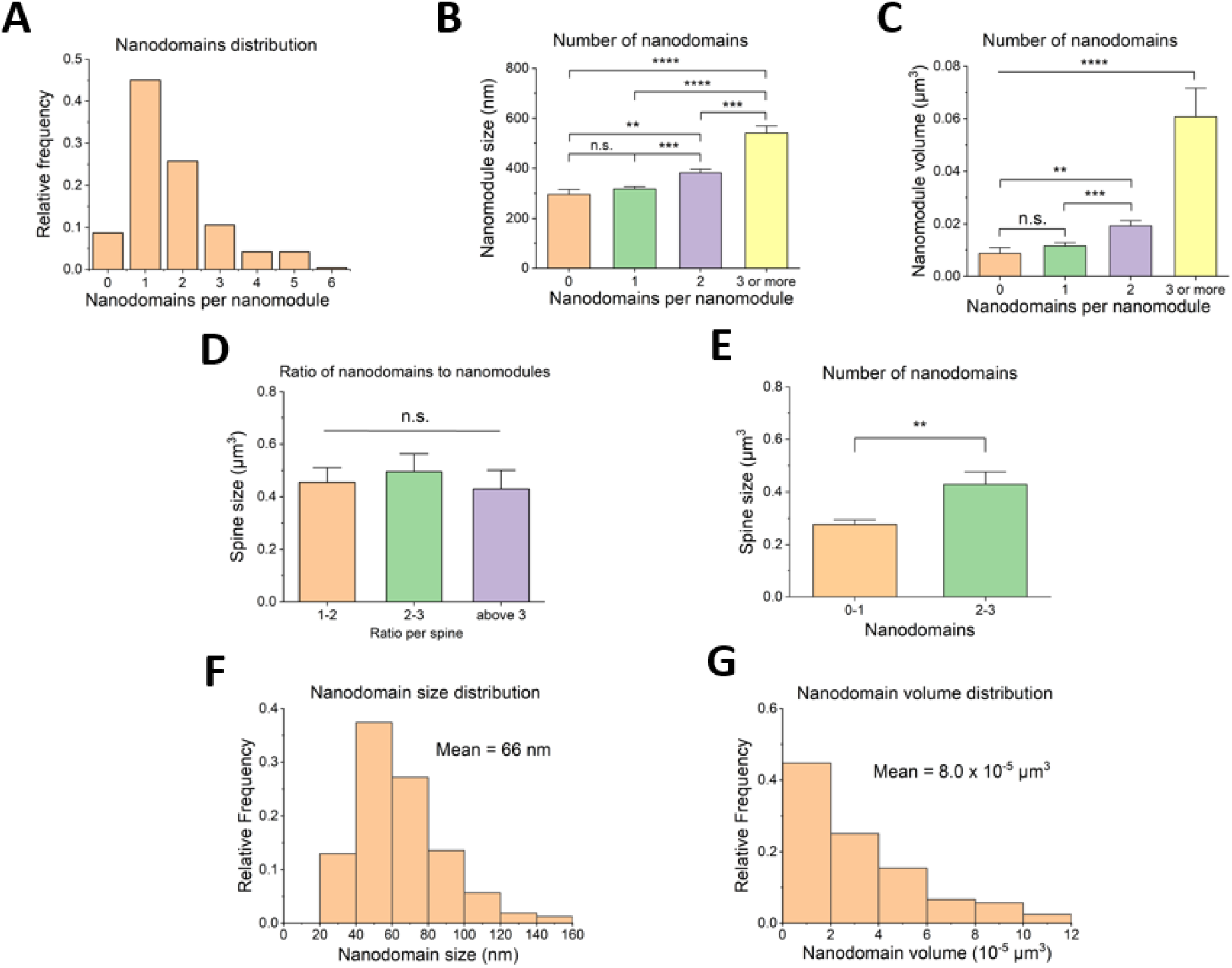
Quantification of nanodomain distribution in spines. (A) Frequency distribution of the number of nanodomains in spines. (B) Nanomodule size versus the number of nanodomains in the nanomodule. (C) Nanomodule volume versus the number of nanodomains in the nanomodule. (D) Spine size vs ratio of the number of nanodomains to the number of nanomodules in the spine. (E) Frequency distribution of the nanodomain size. (F) Frequency distribution of nanodomain volume.

Significance values are determined by Kruskal–Wallis ANOVA. ns : p > 0.05, * : p < 0.05, ** : p < 0.01, *** : p < 0.001. Data is obtained from 4 independent experiments (from 3 mice), dendritic segments of 12 neurons, and a total of 381 spines. Error bars represent standard error of the mean.

## DISCUSSION

In this work, we created a pipeline named ISEPLA for the combined analysis of dendritic spine structure and SMLM localizations of proteins. We applied ISEPLA to identify and quantify nanostructures of synaptic proteins inside dendritic spines. Experimentally, we first took a regular diffraction-limited z-stack of the neuron dendritic segment, followed by SMLM imaging of the target synaptic protein. Computationally, ISEPLA first automatically segments dendritic spines from the diffraction-limited z-stack in 3D. Next, ISEPLA assigns SMLM localizations into clusters, which are distributed onto the segmented spine structures. This allowed us to visualize the nanoscale organization of synaptic proteins in individual spines. Afterward, we used the outputs of this pipeline to quantify the distribution of the synaptic nanostructures inside spines.

Unlike current spine segmentation methods, which are designed primarily for confocal or two-photon microscopy, our approach with DeepD3+Stardist in ISEPLA can perform well under low SNR and resolution conditions typical of widefield imaging in thick brain slice. Despite the specific use of our segmentation approach for correlation with SMLM localization data in our work, it may also be extended to other challenging imaging contexts to enable automatic spine segmentation. The strategy of separating prediction and follow-up segmentation into tasks solved by two different machine-learning models offers a framework that may be applied more generally.

In quantifying spine structures, we focused on spine volume as the sole morphological parameter, rather than classifying spines into traditional categories like mushroom, thin, or stubby. This was firstly due to imaging limitations in thick tissue, where limited resolution and SBR prevent reliable resolution of spine necks, a common pitfall in classifying spines using light microscopy rather than EM images.^46^ Similarly, filopodia—immature dendritic protrusions that lack PSD-95 to form functional synapses—cannot be reliably determined in the images and are therefore excluded from the analysis. Secondly, there is also growing evidence that spine shapes form a continuum without clear categorical boundaries.^47^ On the other hand, it has been shown that the PSD area in synapses correlates strongly with spine head volume but weakly with spine neck parameters.^48^ Thus, we chose to examine the synaptic nanostructure distribution as a function of spine volume. Nevertheless, incorporating additional morphological information may increase the diversity of spine structure representations. In imaging conditions with greater SBR and resolution, where spine necks can be visibly observed, one may extend this method to calculate these parameters from the segmentations. Then, ISEPLA analysis could yield a possibly different pattern of synaptic nanostructure variation with these new parameters. For example, the spine neck morphology may reflect the future stability of the synapse, while spine volume is a direct indicator of current synaptic strength. This could potentially correspond to larger individual nanomodules versus more nanomodules, respectively, in ISEPLA analysis.

To validate our approach to segment spines from diffraction-limited images, we compared the results with super-resolution images, including STORM. When performing STORM imaging of synaptic proteins, dendritic spines are segmented from diffraction-limited images instead of STORM images. The reason that only synaptic proteins but not dendritic segments are imaged by STORM is the result of both experimental and computational constraints. For STORM or other SMLM imaging in thick brain slices, the multiplexing by spectrally separable dyes is limited. Generally, only red-channel dyes are suitable for STORM imaging in thick tissue due to better tissue penetration and chemical properties of the dyes. As a result, only two targets can be reliably imaged in STORM by spectrally separated dyes (e.g., pre- and post-synaptic proteins). However, the rest of the color channels can be utilized for non-STORM imaging targets, as is the case with YFP in our experiments. In addition to this experimental constraint, manually segmenting spines from the STORM or other SMLM methods reconstruction of every FOV is too labor-intensive for the quantitative analysis established in this work. Hence, spine segmentation from diffraction-limited images is a crucial step for correlation with protein localization data, which is robustly established in the ISEPLA method.

After the spine segmentation step, we integrated it with the analysis of SMLM localizations of the synaptic protein PSD-95 to establish ISEPLA. Upon visualizing the localizations of PSD-95 around individual spines, we identify a hierarchical synaptic organization where a spine can contain multiple clusters of PSD-95. These clusters have heterogeneous densities and are themselves made of multiple smaller clusters. The size and shape of the larger clusters match those of “synaptic nanomodules” previously reported in STED studies.^38^ The smaller clusters, on the other hand, match the synaptic nanodomains frequently focused on by SMLM studies.^40,41^ Because past STED studies generally lacked the exquisite resolution of SMLM to resolve nanodomains, it is foreseeable that the previously reported nanomodules are different from nanodomains. At the same time, previous SMLM studies were largely conducted in neuronal cultures that likely did not tend to contain spines with multiple synaptic nanomodules due to a lack of 3D organization, cell density, or more mature spines. While SMLM studies are now turning to imaging in brain slices, the limit with color-multiplexing likely hinders the analysis of synaptic protein distribution inside dendritic spines to study the nanomodule distribution.^49^ These limitations may explain why this relationship between nanomodules and nanodomains has been left unresolved, with the two terms sometimes being addressed as the same.^50,51^. In this study, we addressed these limitations using ISEPLA, which integrates high-resolution SMLM localizations of synaptic proteins with spine segmentation derived from diffraction-limited imaging in a separate color channel, enabling a comprehensive analysis of synaptic architecture within intact tissue.

According to our statistics, two-thirds of the spines do not contain any nanomodules, while it is known that 20% of spines (not including filopodia) lack PSD-95.^52–54^ Some fraction of the other 80% of spines may exhibit PSD-95 puncta in diffraction-limited images, but these puncta do not reach the density requirement defined for nanomodules when examined by ultra-resolution SMLM imaging. Among spines with nanomodules, our results showed that larger spines tend to contain more, but not larger, nanomodules. In turn, larger nanomodules also tend to contain more, but not larger, nanodomains. These observations are consistent with a model in which synaptic growth or strengthening may involve the modular accumulation of substructures—nanodomains within nanomodules, and nanomodules within spines—rather than a uniform scaling of size. As both nanomodules and nanodomains have separately been shown to be connected to synaptic plasticity,^52–54^ this hierarchical organization between them adds another layer of complexity to our understanding of synaptic regulation and functioning.

While we chose to image PSD-95 in this work, another target of particular interest is AMPA receptors (AMPARs) due to their crucial role in synaptic plasticity and alteration in multiple neurological diseases.^57–59^ Recently, the combination of a novel chemical probe and super-resolution STORM has enabled the observation of the nanoscale distribution of surface-bound AMPARs. Our approach is being applied to study the correlation of synaptic versus extra-synaptic AMPARs with the PSD-95 nanostructures in different brain regions and pathological conditions.^60,61^ Finally, applying ISEPLA to models of neurodevelopmental and neurodegenerative disorders may help uncover early alterations in synaptic nano-organization, such as reduced nanomodule density or changes in nanodomain architecture, which could precede overt structural degeneration.

## ACKNOWLEDGEMENTS

The work is funded by the National Institutes of Health grants RF1AG083625 and R01NS100019 (to P.R.S. and H.J.C.). We thank Yongjae Lee (UIUC) from the Selvin lab for nanobody-dye conjugation. We thank Gregory Tracy (UIUC) and Ki H. Lim (UIUC) from the Chung lab for their help in maintaining the mice.

## AUTHOR CONTRIBUTIONS

J.Z., R.V., and P.R.S. designed the experiments. J.Z., R.V., and R.P. performed the experiments.

J.Z. performed data analysis. J.Z. and P.R.S. prepared the manuscript. H.J.C. maintained the IACUC protocol and organized the maintenance of the mouse line. All authors participated in the manuscript revision.

## DECLARATION OF INTERESTS

The authors declare no competing interests.

## DATA AVAILABILITY

Diffraction-limited z-stack images, STORM localizations files after INSPR analysis, analysis codes, and images used in producing the figures are available at: https://data.mendeley.com/preview/wb6c52m7zb.

All raw STORM imaging data, as well as any additional information, will be freely available to those who request and provide a feasible data transfer mechanism (such as physical hard disk drives or cloud storage).

## Notes

### Competing Interest Statement

The authors have declared no competing interest.

### Summary of Updates

Rewriting and additional experimental data and analysis.

