## Supplementary Figure 1 for "Revealing synaptic nanostructure distribution through automatic dendritic spine segmentation and single-molecule localization microscopy"

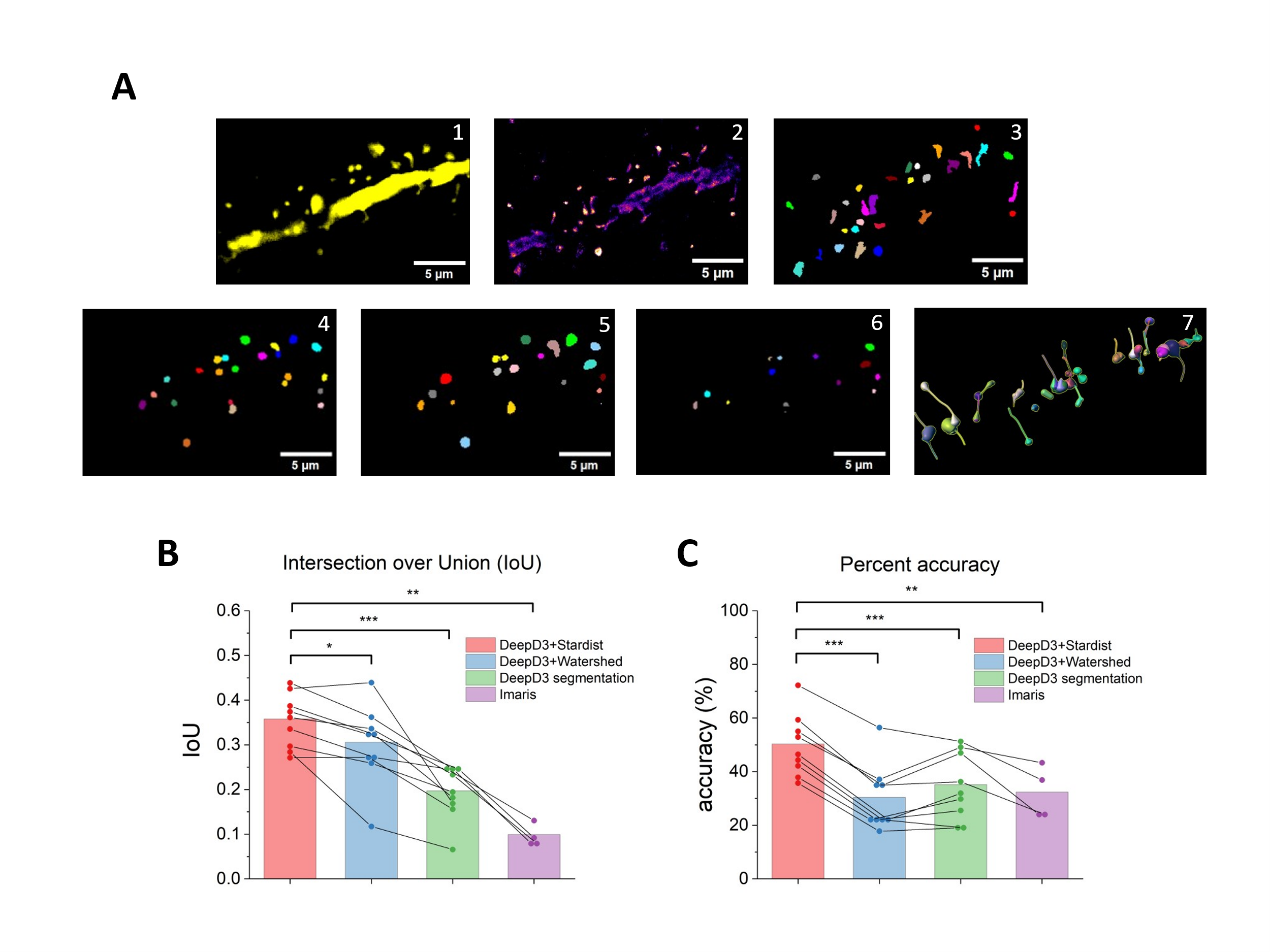

**Supplementary Figure 1. Validation of segmentation methods with super-resolution images**

(A) Fluorescent images of dendritic spines and results of different segmentation methods. 1, One frame of the diffraction-limited fluorescent z-stack after background subtraction. 2, super-resolution reconstruction (STORM or SIM, STORM in this example). 3, Manual segmentation of STORM reconstruction. 4, Segmentation by DeepD3+Stardist. 5, Segmentation by DeepD3+Watershed. 6, Segmentation by the DeepD3 program. 7, Segmentation by Imaris.

(B) Comparison of IoU for different methods. The data points in each method category represent values for one different FOV. Significance values are obtained from paired sample t-tests of DeepD3+Stardist against other methods (alternative hypothesis: mean 1 > mean 2).

(C) Comparison of percent accuracy in the spines correctly detected. Significance values are obtained from paired sample t-tests of DeepD3+Stardist against other methods (alternative hypothesis: mean 1 > mean 2). Chi-squared tests for the contingency table on the total number of spines correctly detected versus incorrectly detected from the four FOVs combined of DeepD3+Stardist against other methods, similarly all give p < 0.01 (**).

* : p < 0.05, ** : p < 0.01, *** : p < 0.001

Data is obtained from 4 FOVs for STORM (89 spines) and 5 FOVs for SIM (144 spines)
